# Genomic description of critical upstream cannabinoid biosynthesis genes

**DOI:** 10.1101/2022.12.15.520586

**Authors:** Peter A. Innes, Daniela Vergara

**Affiliations:** Department of Ecology and Evolutionary Biology, University of Colorado Boulder, Boulder, Colorado

**Keywords:** biochemical pathway, cannabinoids, gene copy number, hemp, marijuana

## Abstract

Cannabinoid production is one of the key attributes of the plant *Cannabis sativa* and the characterization of the genes involved is an essential first step to develop tools for their optimization. We used bioinformatic approaches to annotate and explore variation in the coding genes for critical enzymes comprising the cannabinoid pathway: Olivetol Synthase (OLS), Olivetolic Acid Cyclase (OAC), and Cannabigerolic Acid Synthase (CBGAS), in multiple *C. sativa* genomes. These upstream genes of the Cannabinoid Oxidocyclase Genes THCAS, CBDAS, and CBCAS generate the necessary precursor molecules to produce the cannabinoids THC and CBD. We found that these genes vary in copy number and confirm that OLS, OAC, CBGAS, and the Cannabinoid Oxidocyclases are on separate chromosomes, while homologs are found in proximity. CBGAS, located on Chromosome X, suggests potential dosage effects in female plants. Except for the Cannabinoid Oxidocyclase genes, the other genes have multiple exons, up to 10 in CBGAS. Through differential exon usage explorations in CBGAS we found evidence for potential regulatory differences. This study provides valuable insight on the genomic identity and variation of cannabinoid biosynthesis genes that will benefit future research on the origin and evolution of this pathway, driver of economic, social, and medicinal value.

## Introduction

*Cannabis sativa* L. (marijuana, hemp) is a flowering plant from the family Cannabacea (Bell et al. 2010).**Error! Bookmark not defined**. *C. sativa* likely diverged from its closest extant relative, hops (*Humulus* spp.), approximately 25-27 million years ago in central Asia (McPartland et al 2019; Ren et al 2021). It has been used by humans for more than 10,000 years, making it one of the oldest domesticated plants (Li 1973; Ren 2021; Russo 2007). It is unclear if distinct wild populations of *C. sativa* still exist, though humans have helped spread the cultivated form across the globe (McPartland et al. 2019; Kovalchuk et al. 2020). Worldwide, this versatile plant has been cultivated for multiple uses. Hemp-type *C. sativa* produces fiber that can be used in paper, rope, or clothing, and grain that can be used to extract oil for cooking, personal hygiene and beauty products, while marijuana- type *C. sativa* produces compounds used for medicinal and recreational purposes (Ahmed et al. 2022).

The *Cannabis* genus is considered to have a single species, *Cannabis sativa* L (Watts 2006), but it comprises at least two marijuana-type lineages and one hemp lineage (Sawler et al. 2015; Lynch et al. 2016; Vergara et al. 2016; Kovalchuk et al. 2020; Vergara et al. 2021). *Cannabis sativa* has a mixed mating system with either dioecious or monoecious (with individuals that produce male and female flowers) lineages. Dioecious lineages have sex chromosomes where females are homogametic (XX), and males are heterogametic (XY). Monoecious lineages appear to have two X chromosomes as well (Peil et al. 2003; Hillig 2005; Punja and Holmes 2020).

*Cannabis sativa* is renowned for its production of secondary metabolites, foremost cannabinoids and terpenes. Cannabinoids are a class of compounds that interact with the endocannabinoid system (Gertsch et al. 2008) and many have medicinal (Russo 2011; Swift et al. 2013; Volkow et al. 2014) or psychoactive (Russo and McPartland 2003; ElSohly and Slade 2005) properties. Two of the most widely known cannabinoids are Δ-9-tetrahydrocannabinolic acid (THCA), and Cannabidiolic acid (CBDA), which are converted to the neutral forms Δ-9-tetrahydrocannabinol (THC) and cannabidiol (CBD), respectively, once heated (Hart et al. 2001).

‘Marijuana-type’ lineages used for human consumption has focused on female domestication, since most of the cannabinoid production is found in the trichomes of female flowers (Sirikantaramas et al. 2005; Gagne et al. 2012). There has been strong human selection against males and monoecious individuals. On the other hand, hemp cultivated for their stalks or seeds for fiber or grain production, respectively, produce lower cannabinoid or terpenes (Schafroth et al. 2021).

### Biochemical pathway

The last four steps of the biochemical pathway (Figure 1) from which the cannabinoids are produced begins with <u>steps 1 and 2:</u> the conversion of the compound hexanoyl-CoA to Olivetolic Acid via the action of two enzymes, Olivetol Synthase (OLS) and Olivetolic Acid Cyclase (OAC). In <u>step 3</u> (figure 1) Olivetolic Acid is then converted to another compound, Cannabigerolic Acid (CBGA), via the enzyme Cannabigerolic Acid Synthase (CBGAS). Cannabigerolic Acid, CBGA, has been referred to as the “mother cannabinoid,” since it is the precursor to the cannabinoids produced in <u>step 4:</u> THCA, CBDA, and a third cannabinoid produced in low quantities, Cannabichromenic Acid (CBCA; Page and Stout 2017; Vergara et al. 2019; van Velzen and Schranz 2020; Smith et al. 2021). The enzymes that produce these final cannabinoids are referred to as cannabinoid oxidocyclases (van Velzen and Schranz 2020) because they oxidize the CBGA compound while adding an additional ring.

**Figure 1.**
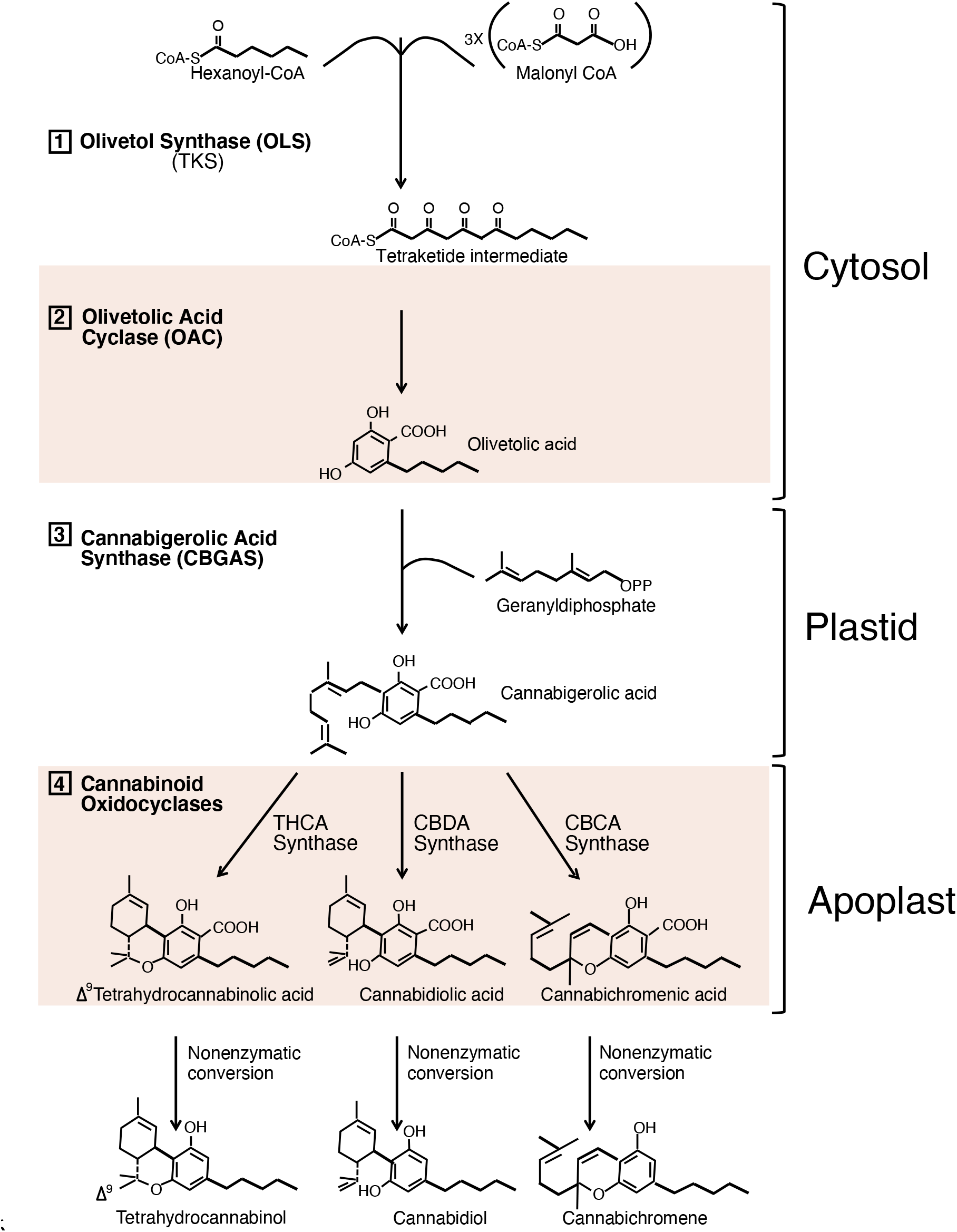
Biochemical Pathway. Last four steps of the biochemical pathway responsible for production of cannabinoids THCA and CBDA. Each step of the pathway has its proposed location in the cell. Figure modified from (Page and Stout 2017; Vergara et al. 2019; Gülck and Møller 2020).

Like THCA and CBDA, decarboxylation is responsible for the formation of Cannabigerol (CBG), Cannabichromene (CBC), and other neutral cannabinoids (Valliere et al. 2019). Due to their low abundance, these “minor cannabinoids” have generally been less well-studied than THC and CBD, although they display a range of interesting pharmacological properties and potential medicinal value (Allen et al. 2022).

### Enzyme-coding genes in the biochemical pathway

Compared to the cannabinoid oxidocyclase genes, which act in the last step of the cannabinoid biosynthesis pathway, less is known about the genes acting upstream: OLS, OAC, and CBGAS. OLS (also known as Tetraketide Synthase, TKS) was cloned and characterized as a plant type III polyketide synthase that produces olivetol (Taura et al. 2009), which is then cyclized by OAC (Gagne et al. 2012) to produce olivetolic acid. OLS and OAC are located on separate chromosomes (Laverty et al. 2019).

The enzymatic activity of CBGAS was described as an aromatic prenyltransferase by which alkylation of olivetolic acid with geranylpyro-phosphase yields cannabigerolic acid (CBGA) (Fellermeier and Zenk 1998). Following an initial genetic sequence description for CBGAS (*CsPT1;* Page and Boubakir 2014) two other similar prenyltransferases (*CsPT4* and *CsPT7*) were described, together forming a cluster (Luo et al. 2019; Rea et al. 2019) that nests within a larger group of aromatic prenyltranserases (Rea et al. 2019; Gülck et al. 2020; Gülck and Møller 2020). Transcriptomic and biochemical analyses suggest that between these three different genes *—CsPT1, CsPT4*, and *CsPT7*— the former two are more likely to produce CBGA *in vivo* (Luo et al. 2019; Rea et al. 2019) as they are highly expressed in glandular trichomes along with OAC and THCAS. On the other hand, *CsPT7* is moderately expressed in trichomes and co-expressed with earlier cannabinoid biosynthesis genes, including OLS (Gülck et al. 2020). Additionally, *CsPT4*, but not *CsPT1*, served as a functional CBGAS in both yeast and tobacco (Luo et al. 2019; Gülck et al. 2020). Whether or precisely how *CsPT1* and *CsPT4* differ in function remains difficult to decipher, as they both have plastid localization signal (Luo et al. 2019; Gülck et al. 2020), and are in close genomic proximity on the X chromosome (Laverty et al. 2019). These findings lead to hypotheses of their possible redundance (Gülck et al. 2020), yet to be experimentally determined. Differences in expression of *CsPT1* have been found between hemp and marijuana (van Bakel et al. 2011), raising questions about how this gene is regulated. Due to the multiexonic structure of the CBGAS genes, alternative splicing may play a role (Laverty et al. 2019).

The diversity, evolutionary origin, function, and genomic context of the Cannabinoid oxidocyclase genes, the last step of the pathway, have been characterized previously (Laverty et al. 2019; van Velzen and Schranz 2020; Grassa et al. 2021). These genes are highly similar at the structural and genetic sequence levels and the enzymes they code for may be classified as “promiscuous” as they act on the same substrate (Onofri et al. 2015; Vergara et al. 2019; Vergara et al. 2020). In their function as synthases, they may also exhibit “sloppiness” (Auldridge et al. 2006; Franco 2011; Chakraborty et al. 2013; Vergara et al. 2020): *in vitro*, they can produce up to eight different products, including each other’s compounds (Zirpel et al. 2018). The genes encoding these synthases are likely derived from the same ancestral gene via duplication events (Onofri et al. 2015; Padgitt-Cobb et al. 2019; van Velzen and Schranz 2020) and are located in close proximity in the genome (Laverty et al. 2019; van Velzen and Schranz 2020; Grassa et al. 2021).

Here we review the genomic locations of the genes coding for the enzymes Olivetol Synthase (OLS), Olivetolic Acid Cyclase (OAC), Cannabigerolic Acid Synthase (CBGAS), and the cannabinoid oxidocyclases (THCAS, CBDAS, CBCAS) and use phylogenetic-based analyses to describe their sequence variation among nine different reference genome assemblies of diverse *C. sativa* lineages. Of these assemblies, four have been assembled to the chromosome level while the remaining five vary in the degree of completion (Gao et al. 2020; Grassa et al. 2021). These upstream genes exhibit duplications and potential copy number variation among assemblies, like the cannabinoid oxidocyclases (Laverty et al. 2019; Vergara et al. 2019; van Velzen and Schranz 2020). Additionally, homologous sequences/copies are in close proximity while the different enzymes are spread throughout the genome on multiple chromosomes (Laverty et al. 2019; Grassa et al. 2021).

We also confirm that these upstream genes have multiple exons, two in the case of OLS and OAC, but up to ten for CBGAS. It is plausible that the multiple exons of the CBGAS-like genes are alternatively spliced, which could influence the genes’ expression or function. Therefore, we examined alternative splicing and found small but significant differences between varieties. Lastly, the location of these CBGAS genes on the X chromosome may have potential implications for cannabinoid production in female plants, which we discuss.

## Methods

### Assemblies

We examined nine different *C. sativa* genome assemblies. These include three hemp types, USO31 (Woods et al. 2021), Carmagnola (Woods et al. 2021), and the previously published Finola assembly (BioProject: PRJNA73819; Laverty et al. 2019). We include the hybrid CBDRx assembly ‘cs10’ (PRJEB29284; Grassa et al. 2021); four marijuana-type assemblies: PBBK (Pineapple Banana Bubba Kush NCBI BioProject PRJNA378470; Vergara et al. 2019), Purple Kush (Purple Kush NCBI BioProject: PRJNA73819) improved from (van Bakel et al. 2011), Jamaican Lion (NCBI BioProject: PRJNA486541) and Cannatonic (NCBI BioProject: PRJNA350523); and the purportedly wild cannabis assembly from Jilong, Tibet (Gao et al. 2020).

### Mining Cannabinoid-related gene sequences

We used BLASTN (Altschul et al. 1990; Gish and States 1993) to find the cannabinoid pathway genes in the aforementioned genome assemblies. For Olivetolic Acid Synthase and Olivetolic Acid Cyclase, steps 1 and 2 on Figure 1, we used the NCBI GenBank cDNA sequences AB164375.1 (Taura et al. 2009), and JN679224.1 (Gagne et al. 2012), respectively.

For Cannabigerolic Acid Synthase which is step 3 on Figure 1, we used previously reported *C. sativa* DNA sequences with putative CBGAS function: CsPT1, CsPT4, and CsPT7 (BK010678, BK010648, BK010683), respectively (Luo et al. 2019). CsPT1 (*C. sativa* prenyl- transferase 1) corresponds to the sequence first reported to be related to CBGAS activity (Page and Boubakir 2014). Initial alignments of these CBGAS-like genes against *C. sativa* assemblies revealed all three have a multi-exonic structure with 10 exons and multiple large introns. To accurately annotate exon boundaries of these genes across assemblies, we searched the cs10 assembly annotation, and found that all three genes were present.

We used the cs10 exons of each gene as BLASTN queries against each of the *C. sativa* genome assemblies. We used the subsequent BLASTN results to extract nucleotide sequences of the exons with BEDTOOLS (Quinlan and Hall 2010), and lastly we manually verified canonical splice site sequences (GT donor and AG acceptor).

For the BLAST queries for the oxidocyclase genes in the last step of the pathway we used the THCAS accessions KP970852.1 (Onofri et al. 2015) and JQ437488.1, the CBDAS accession AB292682.1, and the cDNA for CBCAS (Page and Stout 2017). We used a similar methodology as in (Vergara et al. 2019) where we only recovered sequences with an alignment length of more than 1100 bp.

### ML Trees

We constructed maximum likelihood (ML) gene trees using the program MEGA v.11 (Tamura et al. 2021) for the coding or patial coding regions of OLS, OAC, and CBGAS from each assembly. For all topologies node support was estimated using 500 bootstrap samples and branches with less than 75% support were collapsed. We note that some exons include internal stop codons suggesting they are pseudogenes.

As outgroups, we used BLAST to find similar genes in *Humulus sp*. for both OLS and OAC, which were found for the former but not for the latter gene. For CBGAS, we used two aromatic prenyltransferase genes from *Humulus* (HlPT1-l and HlPT2) as an outgroup, previously shown to cluster with CsPT1, CsPT4, and CsPT7 amidst a broader prenyltransferase gene tree (CITE Gulck et al 2020). Finally, for the cannabinoid oxidocyclase genes we included two cannabinoid-like genes from *Humulus*, 14 sequences from three species from the order Rosales, two of them also from the family Cannabaceae – *Trema orientale* and *Parasponia andersonii* with four and three sequences respectively – and a more distantly related species from the family Moraceae as an outgroup, *Morus notabilis*, with seven sequences.

### Visualization of gene locations and structure, and analysis of alternative splicing

We used the program Chromomap (Anand and Lopez 2020) to map each gene from the cs10 assembly to its respective location. Since CBGA has multiple exons, we used the R package ‘Gviz’ (Hahne and Ivanek 2016) to illustrate the location and size of each exon/intron.

Lastly, for the CBGAS-like genes, we investigated patterns of alternative splicing inferred through differential exon usage within two *C. sativa* varieties, to understand more about their complex multi-exon structure (Supplementary materials).

## Results

### Step 1: Olivetol Synthase

We found multiple copies of OLS-related genes, which include a complete OLS gene, denominated OLS1, as well as a partial OLS gene, which we call OLS2. The nine assemblies differ in copy number of OLS1 (Table S1). Jamaican Lion had four copies, while six other assemblies had two copies, and Cannatonic and PBBK had none. OLS1 is 1,158 bp, with two exons. The first exon is 159bp, the second exon is 999bp, and the intron varies between 103-245bp. OLS2 is in close proximity to OLS1 with 78-80% BLAST percent identity to the second exon of OLS1 and also of approximately 1000 bp. OLS2 is truncated in most assemblies and also varies in copies, with one, two, or up to three copies in Jamaican Lion (table S1).

Although the OLS1 genes from all assemblies cluster together in similarity (Figure 2A), there are two OLS2 genes that cluster with the OLS1 genes instead of the other OLS2 copies. All OLS1 and OLS2 genes are found in chromosome 8 in the cs10 assembly and cs10 has two copies of OLS1-like genes and one copy of OLS2-like genes (Figure 2B). These three genes in cs10 have been annotated, the two copies of OLS1 (LOC115699293 and LOC115700696) and OLS2 (LOC115699292) which was described as a phloroisovalerophenone synthase. The assemblies Finola, Jilong, and Carmagnola have truncated OLS1 genes, and indels causing the premature stop are found within the second exon but all seem to have complete OLS2 genes. We found similar genes in *H. lupulus* that cluster as an outgroup suggesting OLS may be specific to *C. sativa*.

**Figure 2.**
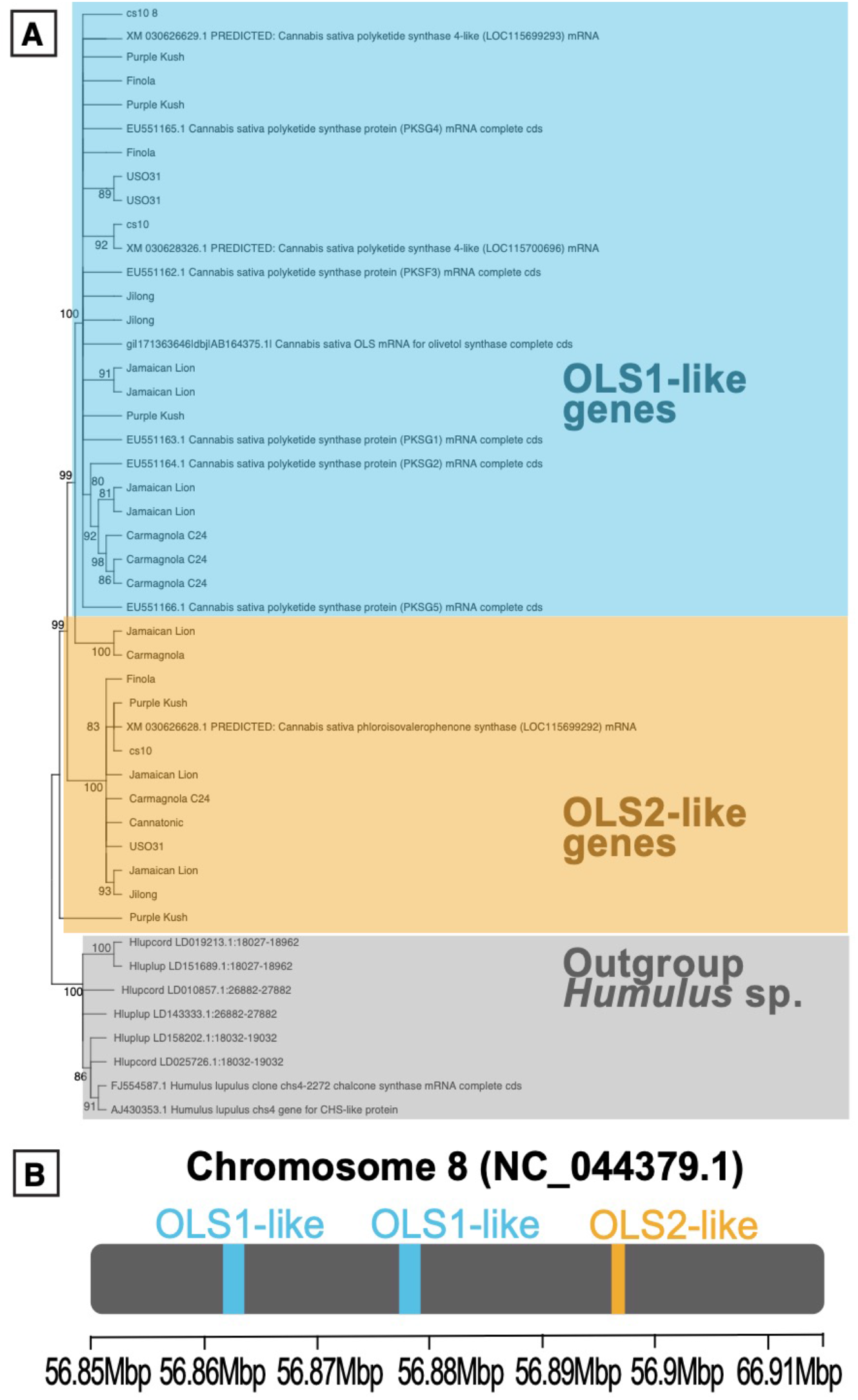
Olivetolic Acid Synthase. **A)** OLS gene tree obtained from seven genome assemblies using Maximum Likelihood approach genome assemblies. OLS was missing from two of the original nine assemblies. Numbers in the branches indicate the percent of 500 bootstraps supporting the topology **B)** Location of the two OLS1 gene copies and the unique OLS2 gene copy in the cs10 assembly.

### Step 2: Olivetolic Acid Cyclase

Like Olivetol Synthase, the nine assemblies differ in number of gene copies of OAC (Table S2), with six copies in Jamaican Lion, four in USO31, three in Carmagnola and PK, two in cs10 and Jilong, one in Finola, and none in Cannatonic and PBBK. OAC also has two exons, the first one of 126bp and the second one of 180bp. The intron is small, approximately 100bp although it varies in the different assemblies (Table S2). USO31 and Purple Kush have truncated genes with indels in the second exon not close to the boundary with the intron. The gene is very similar between varieties/assemblies, and the ML tree reveals substantial polytomy (Figure 3A) that precludes clear relationships between genes from different assemblies. The two OAC copies in the cs10 assembly are found in close proximity in chromosome 9 and have been annotated (LOC115723437 and LOC115723438).

**Figure 3.**
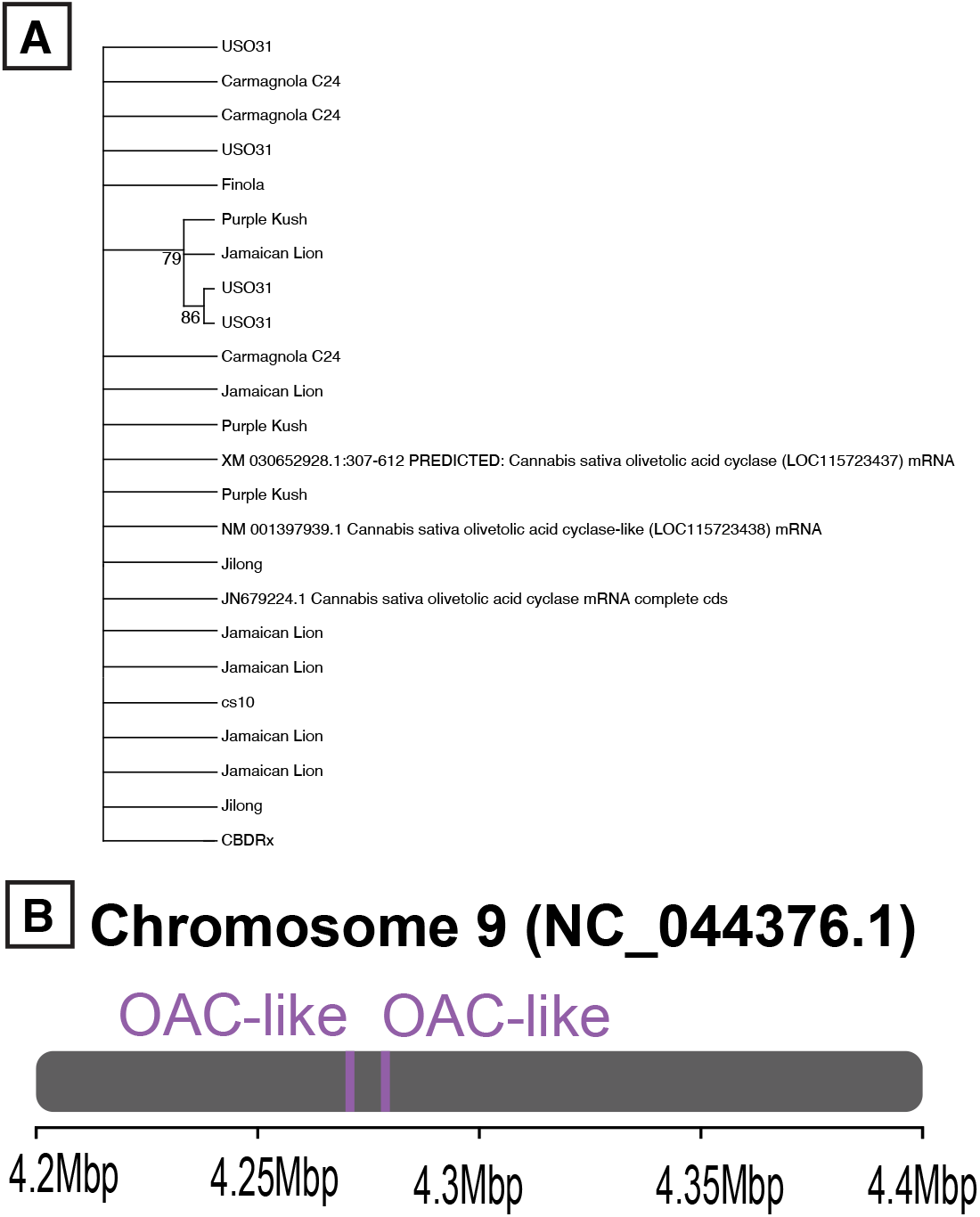
Olivetolic Acid Cyclase. **A)** Relationship via Maximum Likelihood approach between the OAC genes from seven, as it was absent from two of the genome assemblies. Numbers in the branches indicate the percent of 500 bootstraps supporting the topology **B)** Location of the two OAC gene copies in the cs10 assembly.

### Step 3: Cannabigerolic acid synthase

Our initial search for CsPT1, CsPT4, and CsPT7 in the cs10 reference genome annotations revealed near identical matches for all three genes, each with 10 coding exons and annotated as “2-acylphloroglucinol 4-prenyltransferase.” CsPT1 and CsPT7 both had a single top hit (LOC115713215 and LOC115713205, respectively), while CsPT4 closely matched two separately annotated genes: LOC115713171 and LOC115713185. The latter of these two CsPT4 homologs comprised a complete alignment, while the former had a premature stop codon near the end of the seventh exon (our BLAST search identified all 10 exon regions, but only 7 were annotated as ‘CDS’ in the cs10 reference genome annotations). CsPT4 (LOC115713171 but not LOC115713185) and CsPT7 also had multiple isoforms annotated, indicating alternative splicing may play a role in the regulation of these genes. Notably, we found that all CBGAS-like genes are located within the same 193Kb window on the X chromosome (Figure 4).

**Figure 4.**
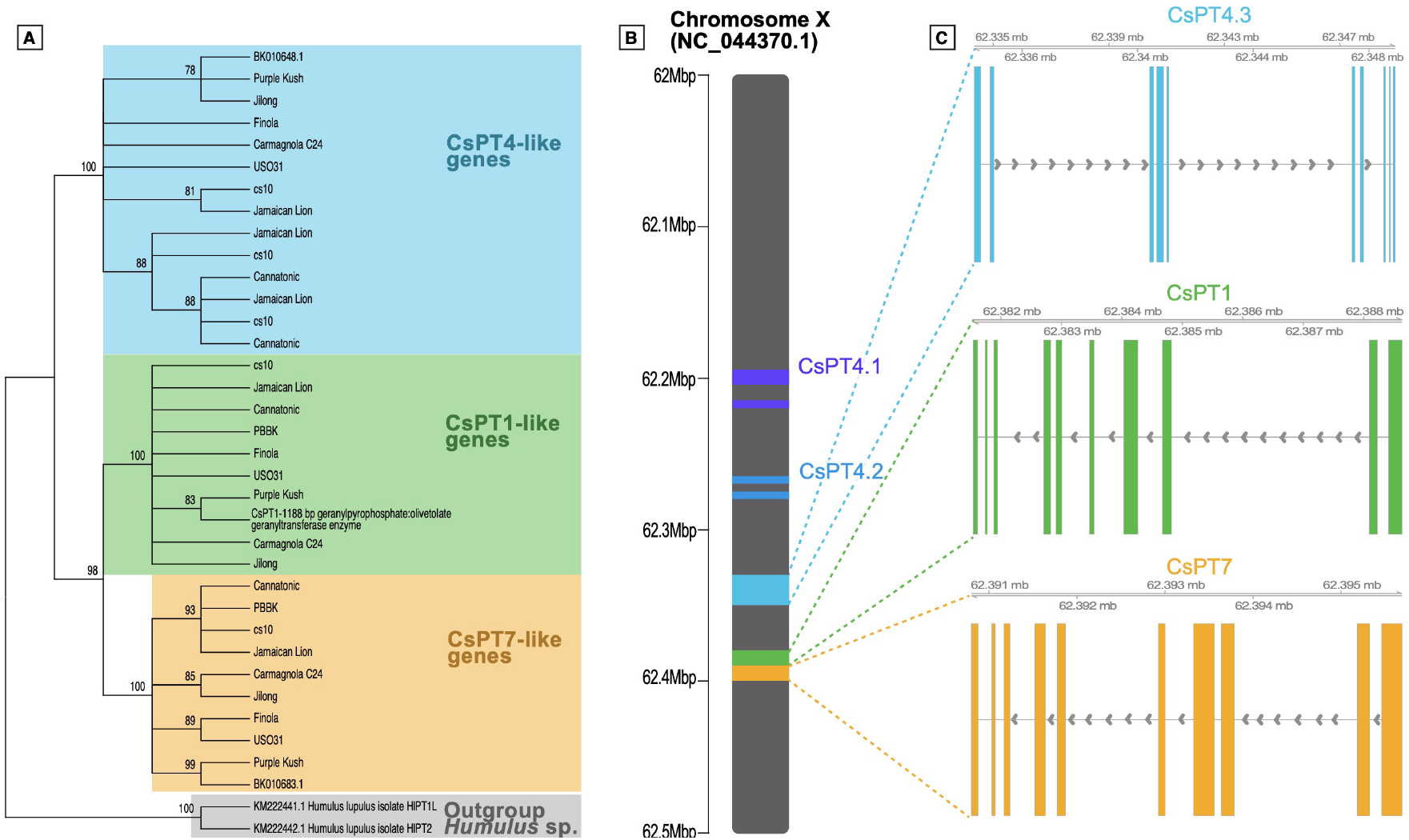
Cannabigerolic Acid Synthase. **A)** Relationship via Maximum Likelihood approach between the CsPT genes from the nine assemblies showing the clear split between CsPT4 from CsPT1 and 7. Numbers in the branches indicate the percent of 500 bootstraps supporting the topology. **B)** Location of the five CsPT gene copies in the cs10 assembly. **C)** Gene structure of three complete copies of CsPT4.3, CsPT1, and CsPT7 showing exons in blue, green, and yellow, respectively. Introns are the uncolored gaps. Arrows indicate the transcription direction.

Subsequent BLAST searches of the CBGAS genes revealed one copy of both CsPT1 and CsPT7 in each of the nine *C. sativa* genome assemblies. Of these, Cannatonic, USO31, and Finola all had internal stop codons in CsPT1; these three assemblies plus PBBK and Carmagnola also had internal stops in CsPT7.

For CsPT4, we found 12 closely homologous sequences across the nine assemblies, indicating potential copy number variation. In cs10, in addition to the two annotated copies of CsPT4, we found a third, previously unannotated copy of this gene. To distinguish these three homologs, we numbered them CsPT4.1-4.3 according to their genomic position in the cs10 assembly. CsPT4.3 corresponds to LOC115713185 in the cs10 reference genome and is a complete ‘canonical’ copy. CsPT4.1 corresponds to LOC115713171, with the premature stop near the end of exon 7, caused by a TGG->TAG point mutation at position 1019. CsPT4.2, the unannotated copy from cs10, was missing likely because it has indel mutations prior to the same TGG->TAG point mutation in exon 7, leading to a more truncated gene. Another distinguishing feature between the three CsPT4 copies is that CsPT4.1 and CsPT4.2 have tandem replications of exon 6, with three and two copies, respectively, although the replications had partial alignments with a total length of 79 out of the total 93 bp of exon 6. Because CsPT4.1 is annotated as a protein coding gene in the cs10 assembly without the additional copy of exon 6, we decided to label the replications of exon 6 as non-coding intronic sequences and did not include them in sequences used for maximum likelihood gene tree analysis.

Among the other assemblies, Cannatonic had CsPT4.1 and CsPT4.2, both with the internal stop codon TGG → TAG at end of exon 7, and Jamaican Lion had three copies (i.e. CsPT4.1, CsPT4.2 and CsPT4.3) only one of which, CsPT4.2, had this same internal stop codon. Purple Kush, Carmagnola, Jilong, and Finola all had a single copy of CsPT4; of these, all were complete except for Finola, which had a 1bp deletion in exon 3 causing an internal stop. Our BLAST search against the USO31 assembly returned hits only for the first two exons of CsPT4. Likewise, PBBK had poor hits for various CsPT4 exons but lacked a complete copy (Table S3).

Phylogenetic-based analysis of the CBGAS gene sequences from all assemblies revealed three well supported clades, corresponding to CsPT1, CsPT4, and CsPT7 (Figure 4). CsPT1 and CsPT7 are more similar to each other than either one is to CsPT4. Within the CsPT4 clade, there is one well-supported split that separates complete/functional from truncated and/or non-functional copies, except for CsPT4_Finola (which has a 1bp deletion in exon 3 causing an internal stop), grouping with the complete/functional CsPT4 copies from other assemblies.

The CBGAS-like genes (CsPT1, CsPT4, CsPT7) contain multiple large introns (Table S3). In cs10, intron 2 of CsPT1 is 3,553bp long. CsPT4 has an even longer second intron compared to CsPT1: in cs10 it is 5,386bp long (CsPT4.3), and even larger in Purple Kush, at 5,946 bp. Intron 5 of CsPT4 is even longer than its intron 2, ranging from 6,344 bp in cs10 to 8541 in Purple Kush, for the ‘canonical’ CsPT4 copies (i.e. those without premature stops codons).

### Differential splicing (exon-usage) analysis in CBGAS

We found evidence for differential splicing of CBGAS-like genes between different varieties. Specifically, we found significant differential exon usage (DEU; false discovery rate < .05) in all four CBGAS-related genes annotated in cs10 (Figure S1) with CsPT1 and CsPT4.3 having the highest overall expression. Notably, CsPT1 showed nearly three-fold higher expression in White Cookies compared to Canna Tsu (Figure S1 A). CsPT1 showed significant DEU for exons 4, 5, 9, and 10. CsPT4.3 showed DEU only for exon 8. CsPT4.1 was the lowest expressed gene and nearly absent in Canna Tsu, though showed substantial DEU (Figure S1 C). Exons 1, 8, 9 and 10 of CsPT7 showed significant differential usage (Figure S1 D).

### Step 4: Cannabinoid oxidocylase genes

We found 143 genes in the nine different assemblies related to the cannabinoid oxidocyclase genes. These same 143 genes were recovered regardless of whether THCAS, CBDAS, or CBCAS was used as the BLAST query. Jamaican Lion was the assembly that contained the greatest number of oxidocyclase genes (32) and Jilong the fewest (8).

Our ML gene tree reconstruction revealed relationships both between different oxidocyclase gene copies and homologs in different assemblies (Figure 5A). We found 44 genes related to CBCAS, seven to THCAS, and 17 to CBDAS. We also found five clusters of genes with unclear function, labeled in Figure 5A as ‘Unknown’ groups 1, 2, and 3, ‘Additional oxidocyclase-like genes’, and ‘Additional uncharacterized genes’. Unknown groups 1 and 2 contains 13 genes, six and seven genes, respectively. Unknown group 3 has 45 genes from all nine assemblies which is 32% of all genes. In other words, each genome assembly contains genes from this unknown 3 cluster. The six genes labeled ‘Additional oxidocyclase-like genes’ form a polytomy alongside the CBDAS and the ‘Unknown 3’ clusters, making it unclear which of these sets of genes is sister to the clade comprising THCAS, CBCAS, ‘Unknown 1’, and ‘Unknown 2’.

**Figure 5.**
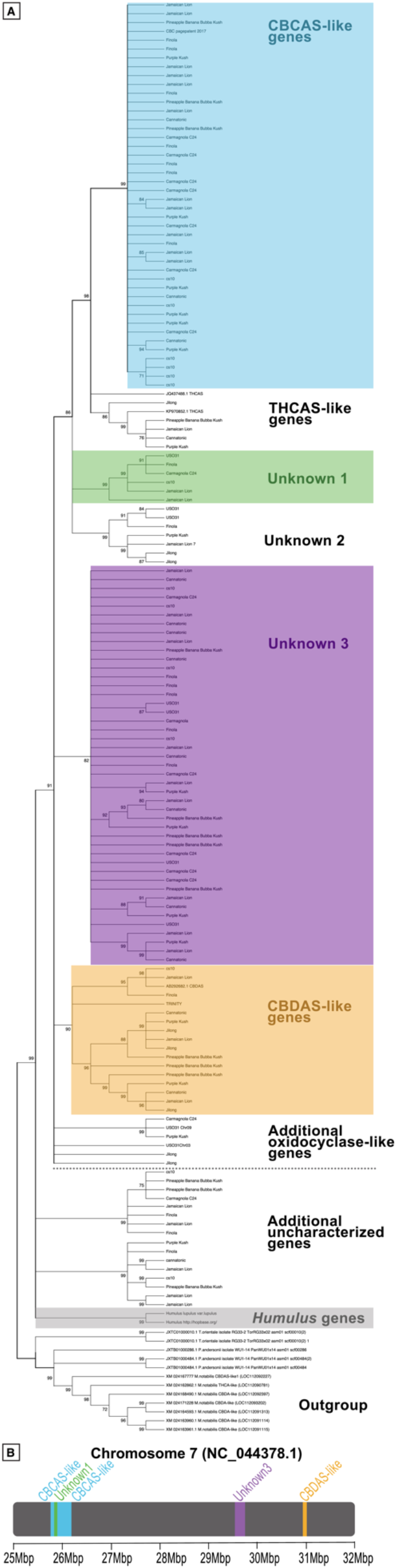
Cannabinoid oxidocyclase genes. **A)** Gene tree of cannabinoid oxidocyclases and related sequences via Maximum Likelihood approach from the nine assemblies, showing the clear split between CBCAS in blue and THCAS uncolored, three unknown clusters in green, uncolored, and purple, respectively, CBDAS in yellow and two groups with ‘additional oxidocyclase-like genes’ and ‘Additional uncharacterized genes, both uncolored. The *Humulus* sp. genes are colored in gray. Numbers in the branches indicate the percent of 500 bootstraps supporting the topology. The dashed line separates the primary cannabinoid oxidocylase clade from more dissimilar sequences **B)** Location of the 12 oxidocyclase genes in the cs10 assembly (tandem repeats are collapsed into a single color block).

The cannabinoid oxidocyclase genes CBCAS and THCAS are very similar and are sister to each other in our gene tree. Every assembly except for USO31 has both complete and truncated CBCAS-like genes. However, only marijuana-type individuals contain THCAS-like genes. CBDAS forms a group with two clusters, one of them with only truncated genes (Figure 5A).

Considered by genome assembly, the Carmagnola assembly lacks THCAS and CBDAS genes but has one complete CBCAS gene, which is the only complete cannabinoid oxidocyclase gene we found from this assembly. USO31 lacks THCAS and CBCAS genes but has complete CBDAS. The last hemp-type genome assembly, Finola, has a complete CBDAS and CBCAS genes, but no THCAS-like genes.

The hybrid individual cs10 does not have THCAS genes but does have complete CBDAS and CBCAS genes.

All marijuana-type genome assemblies contain one THCAS-like gene which is complete for all individuals except Cannatonic. Interestingly, Cannatonic is usually a 1:1 CBD:THC or high CBD variety. The only marijuana-type genome that has a complete/functional CBDAS-like gene is Jamaican Lion.

The ‘Additional uncharacterized genes’ from the different assemblies have a BLAST identity of less than 80% to the oxidocyclase genes THCA, CBDA, and CBCA (Table S4). These cluster into two groups of eight genes for 16 genes total, are found outside the oxidocyclase clade, and have the same relationship to the genes from *Humulus* sp as to the oxidocyclase genes in the ML tree (Figure 5A). All assemblies except for USO31 had one of these genes, some of them complete (Table S4).

All cannabinoid oxidocyclase genes are found in chromosome 7 in the cs10 assembly (Figure 5B), but those ‘Additional uncharacterized genes’ are found in chromosome 2.

## Discussion

These cannabinoid biosynthesis genes described in this study have both societal and legislative implications. The compounds they produce are used for setting legal definitions for different categories of cannabis products, as well as for ‘chemotaxonomic’ purposes to classify different *C. sativa* varieties (Henry et al. 2018; Reimann-Philipp et al. 2019; Smith et al. 2021). For example, the legal definition of hemp in the United States is any *C. sativa* plant containing 0.3% THC or less (Small and Marcus 2003).

We describe these cannabinoid biosynthesis genes and provide new insights about their sequence and potential copy number variation, and genomic location across diverse *C. sativa* lineages and genome assemblies. Our results suggest that independent of the lineage, whether a hemp-type or a marijuana-type *C. sativa*, these genes related to the cannabinoid pathway can be found in multiple copies and are scattered throughout the genome.

### Steps 1 and 2: OLS and OAC

Our results strengthen reports of duplicate copies of OLS in Purple Kush and Finola (van Bakel et al. 2011; Laverty et al. 2019) and of OAC in Tachllta Till, a marijuana-type variety (McGarvey et al. 2020). We find tandem duplications of both these enzyme genes in the genome assemblies of Carmagnola, Finola, cs10, Purple Kush, Jamaican Lion, and Jilong. The fact that both duplications are present in the Jilong assembly, which is a putative wild genotype, suggests the duplication events could have arisen before strong artificial selection pressures from humans. Future research should investigate how prevalent the OLS and OAC duplications are, and whether the gene copies are functionally redundant, or have any impact on gene dosage. Additionally, for OLS, future research could determine whether the OLS2-like gene, similar to the second exon of OLS1 is functional and what products it yields.

### Step 3: CBGAS

We find evidence for potential copy number variation in CBGAS: in the cs10 assembly, arguably the most complete and well-assembled of the nine, the CsPT4 gene is repeated three times in close proximity, although just two of the copies, CsPT4.3 (complete, 10 exons) and CsPT4.1 (truncated, seven exons), are annotated as protein coding genes. Two other assemblies, Jamaican Lion and Cannatonic, also have additional copies of CsPT4. The replications of CsPT4 in cs10 are intriguing and deserve further investigation, for instance, do they exist in other genotypes not included here?

Based on the close homology and proximity of CsPT1, CsPT4, and CsPT7, there has apparently been an extended history of gene duplication at this locus, though the functional consequences, if any, remain unclear. Finola (hemp) and Purple Kush (marijuana-type) do not differ in number of CBGAS (neither CsPT1 or CsPT4) gene copies, yet findings report more than five-fold greater expression of CsPT1 (Aromatic Prenyltransferase, referred by van Bakel as the ‘*AP*’ locus) in Purple Kush compared to Finola (van Bakel et al. 2011). It was later suggested that the presence of repeat sequences and indels within introns of CsPT1 might regulate its transcription or alternative splicing (Laverty et al. 2019). We indeed find evidence for differential splicing of CsPT1 and multiple other CBGAS-related genes (Figure S1) between a high THC variety (White Cookies) and a high CBD variety (Canna Tsu), but with a small magnitude of difference as well as sample size, and therefore should be confirmed by additional studies. However, in support of our findings, a recent study reported alternative splicing events at the CBGAS locus, as well as others in the cannabinoid biosynthesis pathway, though tests for differential splicing between varieties were not performed (Wu et al. 2021).

Previous studies have reported varying lengths of intron 2 in CsPT1 (McGarvey et al. 2020), in agreement with our results. We found that CsPT4 has an even longer second and fifth introns compared to those previously reported for CsPT1. The implications of these very large intronic regions, although unclear, should motivate future studies.

The presence of CBGAS genes (both CsPT1 and CsPT4) on the X Chromosome (Chromosome 10) is also something that has not been investigated closely and has potential implications of higher dosage in females who contain two X chromosomes. Females produce higher concentration of cannabinoids due to the abundance of trichomes on their flowers (Sirikantaramas et al. 2005; Gagne et al. 2012). Perhaps that these genes are found on the X chromosome could be another of the underlying reasons of the sex difference in cannabinoid production.

### Step 4: Cannabinoid Oxidocyclase genes

Our results support previous findings that the cannabinoid oxidocyclase genes are numerous, in close proximity in the genome (Laverty et al. 2019; Grassa et al. 2021), and vary in copies (van Velzen and Schranz 2020; Vergara et al. 2021) between assemblies (Table S4). Through this study we found that THCA is only found in marijuana-types but not in hemp. This raises the question or whether or not these are derived genes only found in some of the *C. sativa* lineages. Although the idea of some of the cannabinoid oxidocyclase genes being ancestral has been proposed before (Onofri et al. 2015), we cannot conclusively support or disprove it given that no other species is known to share this biosynthetic pathway (Vergara et al. 2019). However, the recent sequencing of the putative wild specimen from Jilong, Tibet, with very few cannabinoid oxidocylcase genes, suggests that increased gene duplication events may be associated with domestication (Gao et al 2020).

We also found that the assembly of Cannatonic, which is a variety sold as a 1:1 CBD:THC, has truncated sequences for both the canonical THCA and two CBDA synthase genes, making it unclear which of the other 13 genes (Cannatonic has 15 cannabinoid oxidocyclase genes total) are responsible for the production of CBDA and THCA. Therefore, although we only have one genome assembly for Cannatonic, our results suggest that despite the truncated genes, these compounds may still be produced. If this is the case, and other oxidocyclase genes are producing the THCA and or CBDA compounds, it will make it hard for farmers to produce compliant hemp below the 0.3% THC regulated limit. This result supports the idea that the enzymes codified by these oxidocyclase genes are both promiscuous (Onofri et al. 2015; Vergara et al. 2020) and sloppy (Auldridge et al. 2006; Franco 2011; Chakraborty et al. 2013; Zirpel et al. 2018; Vergara et al. 2020).

The large number of CBCAS-like genes is also an intriguing finding given the low production quantities of this cannabinoid, which is the reason why it is considered a ‘minor’ cannabinoid. However, despite low productions of CBCA, expression levels of this gene varies between *C. sativa* varieties (Allen et al. 2022). As previously suggested, perhaps CBCAS is the sloppiest of all synthases in the pathway (Vergara et al. 2020). Similarly, the genes whose products remain to be established are interesting due to their large number. These two results regarding the numerous CBCAS-like genes despite low production of CBCA, and those with still unknown products support the notion that all these genes are acting in unison as a network to produce the complex cannabinoid phenotype (Zager et al. 2019). Our results also suggest that the genes whose products remain unknown are either producing one of the three known cannabinoids THCA, CBDA, and or CBCA, or that they have not been directly associated to the cannabinoid compound—from one of the numerous cannabinoids found in the plant—they are producing, or that there may still be cannabinoid compounds yet to be discovered.

With different phylogenetics-based approaches, the relationship between the oxidocyclase genes remain the same as previously determined (van Velzen and Schranz 2020): CBCAS and THCAS are sister clusters to each other, various clusters of genes with unknown function exist, and some genes that fall outside the oxidocyclase-like clade that could be closer to *Humulus* sp. genes (van Velzen and Schranz 2020).

### Entire pathway

All the enzymes from different steps of the pathway are found in different chromosomes, far apart from each other. This suggests that through recombination, individuals may frequently inherit novel combinations of cannabinoid biosynthesis-related genes and alleles that will work together to produce the cannabinoid phenotype. There may be particular alleles of each gene, or combinations of genes in the entire pathway that, when together, could enhance or diminish the production of these compounds. Because these genes are unlinked, this could facilitate artificial selection in eventually combining the best performing genes and alleles throughout the entire pathway to enhance cannabinoid production.

Through our phylogenetic-based gene tree analyses, we found that some of these genes harbor more sequence variation across different *C. sativa* varieties than others. OAC has the least amount of sequence variation, producing a large polytomy in the gene tree, while the CBGAS-like genes and the cannabinoid oxidocyclases show more distinct clustering and branching, suggestive of sequence divergence.

Given that the cannabinoid compounds discussed here are unique to *C. sativa*, our findings, in agreement with previous studies (van Velzen and Schranz 2020), suggest that the enzymes producing these compounds acquired a new function after the speciation event with *Humulus* species approximately 25 mya (Bell et al. 2010). Because all of the genes in the pathway (OLS, OAC, CBGAS, and the oxidocyclases) appear to have a significant history of duplication, each of these genes could be going through their own process of either concerted evolution, in which the copies maintain similar sequence and function, neofunctionalization, where one of the copies acquires a novel function, or subfunctionalization, where the original function of the gene becomes split among the copies (Lynch 2007; Zhang 2013). The apparently major role of gene duplication in the *C. sativa* cannabinoid biosynthesis pathway is consistent with the evolution of plant secondary metabolism pathways more broadly (Ober 2005; Hartmann 2007).

The lack of clustering of cannabinoid biosynthesis genes in the *C. sativa* genome is similar to the genomic organization of some other plant specialized metabolism pathways, in which genes are spread across multiple chromosomes, for example anthocyanin and saponin production in carrot (Yildiz et al. 2013) and *Nicotiana benthamiana* (Khakimov et al. 2015), respectively, and glucosynilate biosynthesis in *Arabidopsis thaliana* (Kliebenstein and Osbourn 2012; Hofberger et al. 2013). However, a growing number of studies have reported the opposite pattern, with genomic clustering of genes that interact in the same secondary metabolism pathway (Nützmann and Osbourn 2014). Whether or not cannabinoid biosynthesis pathways in other species (e.g. *Radula marginata, Rhododendron dauricum*) form genomic clusters is unknown, due to lack of genomic resources (Taura et al. 2016; Hussain et al. 2018; Saeki et al. 2018; Hussain et al. 2019). The evolutionary consequences of having linked versus unlinked represent an intriguing avenue of future comparative genomics research, especially in the context of *C. sativa* crop improvement. Independent of the location of the genes in the cannabinoid pathway, our study provides insights on the future possibilities of modifying the genes upstream in the pathway to silence the entire cannabinoid production downstream, which would be useful for hemp cultivators to assure the maintenance of plants with less than 0.3% THC.

### Caveats

Our study comes with limitations, one is the sample size and the other one is the genome assembly quality. Although the individuals included in this study comprise the variation found in *C. sativa* as they are part of different lineages within the species, these are only nine genome assemblies. Therefore, all the results reported here belong to a small subsample within the species.

Our survey highlights the difference in assembly quality among currently available *C. sativa* genomes. The known genome size of *C. sativa* is approximately 800Mb, and the cs10 assembly is currently the representative genome on NCBI GenBank (Grassa et al. 2021). However, among the included genome assemblies in our study, size varied from ∼500Mb (PBBK and Cannatonic) to ∼1Gb (Jamaicon Lion). Given these substantial size differences, it is likely that the PBBK and Cannatonic assemblies are both missing genomic regions and that the Jamaican Lion assembly has redundancies. Indeed, we found that PBBK and Cannatonic lacked crucial genes (e.g. OLS, OAC), while Jamaican Lion often had more gene copies compared to other assemblies. Thus, the putative copy number variation among PBBK, Cannatonic, and Jamaican Lion compared to other assemblies could be attributed to sequencing or assembly issues rather than real biological copy number variation. However, for Jamaican Lion, the abundance of additional cannabinoid biosynthesis gene copies might represent important allelic variation and should not be completely disregarded. Due to this potential allelic diversity from the Jamaican Lion assembly, and the fact that we cannot distinguish between alleles or gene copies, we decided to retain all sequences from this assembly in our gene trees. All analyzed assemblies in our study provide valuable resources for *C. sativa* genomic research, but future studies should carefully consider differences in assembly completeness if making comparisons between varieties/assemblies.

## Acknowledgments

The authors would like to thank the graduate and undergraduate students of the 2020 Comparative Cannabis Genomics course at the University of Colorado, Boulder, from which this study developed. We also thank Nolan Kane, Rick E. Miller, Liam Friar, and other Kane Lab members for helpful comments and discussion of earlier versions of the manuscript.

## Author contributions

PAI and DV performed all analyses and wrote the manuscript.

## Competing financial interests

DV is the founder and president of the non-profit Agricultural Genomics Foundation and the sole owner of the company CGRI, LLC.

## Supplemental materials

Analysis of differential exon usage in CBGAS genes between two *C. sativa* varieties

## Methods

### CBGAS-like genes differential expression

We downloaded publicly available transcriptomic data from floral tissue of two *C. sativa* varieties, White Cookies (high THC) and Canna Tsu (high CBD), which were grown in a common environment (Zager et al. 2019). We aligned three RNA-seq libraries from both varieties to the cs10 reference genome (Grassa et al. 2021) using STAR in two-pass mode (Dobin et al. 2013). We then counted reads at the (exon) level using featureCounts (Liao et al. 2014) of only uniquely mapping reads. We tested for differences in relative exon usage between varieties for CsPT1, CsPT4.3, CsPT4.1, and CsPT7, using DEXSeq (Anders et al. 2012). Due to the absence of CsPT4.2 in the cs10 annotation, we did not include this gene in our differential exon usage analysis. The used approach controls for differences in overall gene expression level between conditions and is an indicator of certain types of alternative splicing, for instance exon skipping or mutually exclusive exons.

**Figure S1.**
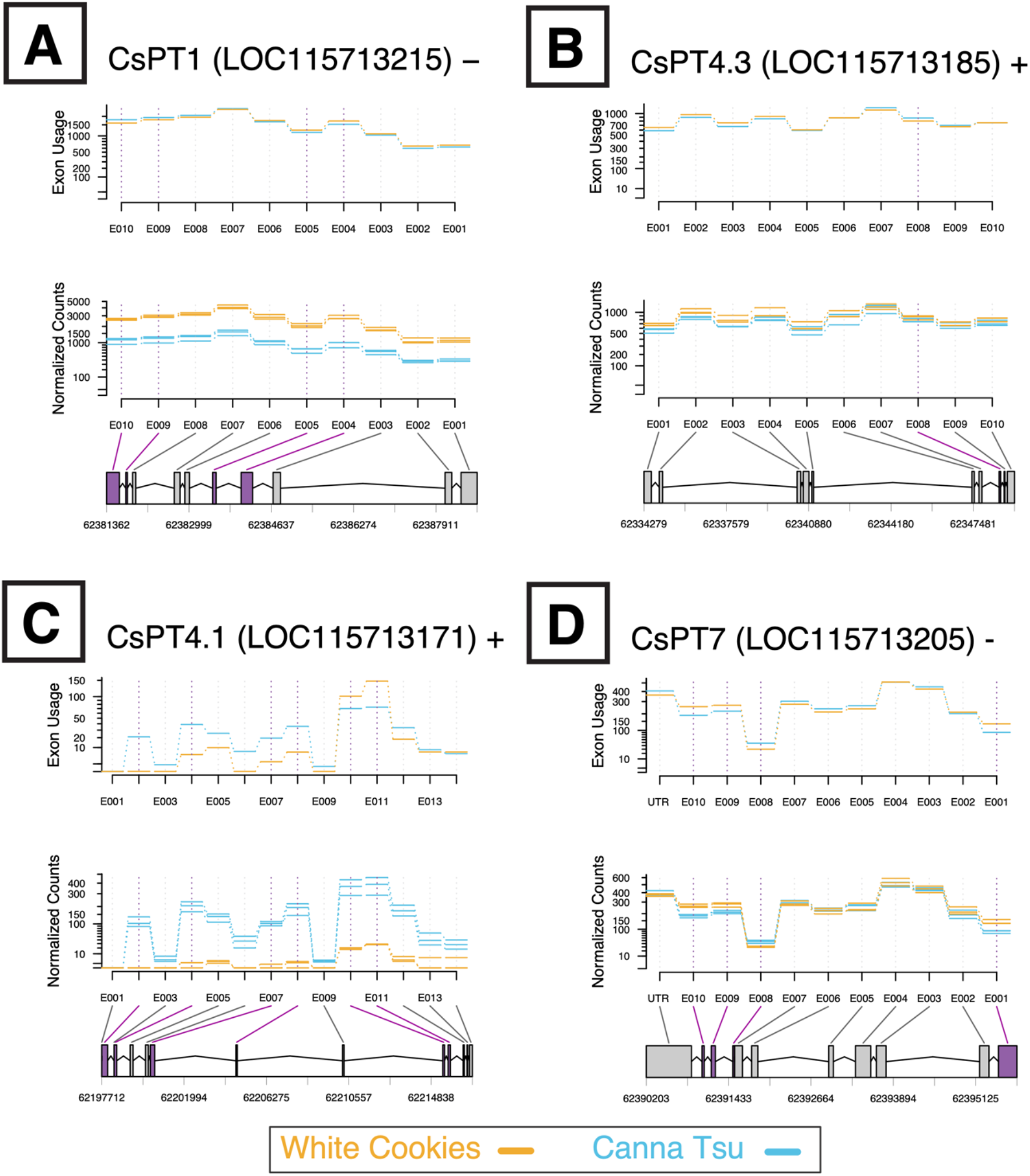
Differential exon usage diagrams for different CBGAS-like genes. The high-THC variety ‘White Cookies’ is shown in yellow, and the high-CBD variety ‘Canna Tsu’ is blue. Exons shaded in purple have significant differences in relative expression (i.e. ‘usage’) between varieties (FDR < .05), independent of any gene-level differences in expression. Normalized read counts for each sample are shown in the middle section, while bottom sections show gene structure and location on chromosome X (NC_044370.1) of four of the five CBDAS-like genes from the cs10 reference assembly: **A)** CsPT1; **B)** CsPT4.3; **C)** CsPT4.1; and **D)** CsPT7

